# Plant species and soil type influence rhizosphere bacterial composition and seedling establishment on serpentine soils

**DOI:** 10.1101/489344

**Authors:** Alexandria N. Igwe, Rachel L. Vannette

## Abstract

Root-associated microbial communities influence plant phenotype, growth and local abundance, yet the factors that structure these microbial communities are still poorly understood. California landscapes contain serpentine soils, which are nutrient-poor and high in heavy metals, and distinct from neighboring soils. Here, we surveyed the rhizoplane of serpentine-indifferent plants species growing on serpentine and non-serpentine soils to determine the relative influence of plant identity and soil chemistry on rhizoplane microbial community structure using 16S rRNA metabarcoding. Additionally, we experimentally examined if locally adapted microorganisms enhance plant growth in serpentine soil. Plant species, soil chemistry, and the interaction between them were important in structuring rhizoplane bacterial communities in both the field and experimental soils. In the experiment, rhizoplane microbial community source influenced seedling survival, but plant growth phenotypes measured were largely invariant to microbial community with a few exceptions. Results from the field sampling suggest that plant species associate with specific microbial communities even across chemically distinct soils, and that microbial communities can differentially influence seedling survival on harsh serpentine soils.

## Importance

Microbial communities on plant roots can influence host plant phenotype, survival, and fitness, with community and ecosystem consequences. However, which factors structure microbial community structure is poorly understood, particularly across large gradients in soil properties. The survey and experiment described here shows that plant species host distinct microbiomes despite strong turnover in soil chemistry, although soil type also influences bacterial structure. These results suggest that host specificity and soil conditions are both strong drivers of rhizoplane bacterial composition in serpentine ecosystems and influence plant seedling establishment. These results imply that plant species can recruit relatively convergent microbiomes from distinct microbial communities found on different soil types and that soil microbial communities influence seedling establishment.

## Introduction

Root-associated microbial communities are important mediators of plant traits and soil processes. For example, microorganisms can accelerate nutrient cycling and increase nutrient availability to plants (Baker *et al*. 2018). Some microbes fix nitrogen (Boyd and Peters 2013; Mus *et al*. 2016) and provide plants with nutrients (Shakeel *et al*. 2015), with effects that scale to influence global nutrient cycles (Finzi *et al*. 2015). Further, root-associated microorganisms can alter plant traits such as disease tolerance (Santhanam *et al*. 2015), root architecture (Zhou *et al*. 2016), and drought tolerance (Lau and Lennon 2011). However, the factors that contribute to the structure of root-associated microbial communities are diverse and their relative influence across scales are poorly understood.

Both plant identity and soil chemistry influence root microbiome composition (Burns *et al*. 2015; Erlandson *et al*. 2018; Leff *et al*. 2018), but their relative influence remains difficult to estimate. Plants can influence root-associated microbial communities by exuding carbon-rich compounds from their roots, which can select for beneficial plant-growth-promoting bacteria (Badri and Vivanco 2008; Chaparro *et al*. 2013) and in some cases, can feed back to increase plant biomass (Lally *et al*. 2017). Root-associated microbes can contribute to environmental tolerance of plant hosts via processes such as phosphorus solubilization (Govindasamy *et al*. 2010; Khan, Zaidi and Ahmad 2014), production of extracellular polymeric substances (EPS) (Glick 2012), and metal complexation (Tak, Ahmad and Babalola 2012). Although most work to date has examined soil or rhizosphere microbiome composition, rhizoplane communities can provide protection against soil-borne pathogens (Deora *et al*. 2005; Islam *et al*. 2005) and are an important barrier between rhizosphere and endosphere communities (Edwards *et al*. 2015). Additionally, microbes in the rhizoplane may be more closely linked to plant benefits, including drought tolerance (Fitzpatrick *et al*. 2018) than microbes found in the rhizosphere. On the other hand, soil physical and chemical properties can also be strong drivers of rhizosphere and rhizoplane microbial community composition, particularly on large scales or across soils that vary widely in composition. For example, pH (Bartram *et al*. 2014; Wu *et al*. 2017), salinity (Sardinha *et al*. 2003; Dillon *et al*. 2013), drought (Barnard, Osborne and Firestone 2013; Chodak *et al*. 2015), and the presence of heavy metals (Wood *et al*. 2016) all influence soil and root-associated microbial community structure. Still, the relative importance of plant identity and soil chemistry across soil gradients remains difficult to predict in large part because plant species are often restricted to particular soil types.

Both soil and plant identity can shape not only structure but also function of rhizoplane microbial communities. Soil stressors, including drought, can promote the development of locally adapted microorganisms (Hawkes and Keitt 2015) which have been shown to be important for plant growth (Lau and Lennon 2012; Revillini, Gehring and Johnson 2016). On the other hand, plant-induced changes in soil parameters, including microbial communities, can feed back to impact plant growth and microbial communities (Kulmatiski *et al*. 2008; Rúa *et al*. 2016). By associating with microbes from a shared soil environment, plants are sometimes able to gain a greater fitness advantage than if they associated with microbes from a different soil environment (Rúa *et al*. 2016; Gehring *et al*. 2017). However, in some cases, association with sympatric microorganisms results in no significant changes in fitness relative to uninoculated plants (Doherty, Ji and Casper 2008). Overall, there is still much to be learned about the role of locally adapted microorganisms for plant growth, particularly if, when and how they may influence plant hosts.

Serpentine soils are an ideal system in which to examine local adaptation and the relative contribution of plant identity and soil chemistry to microbial community structure. Serpentine soils are characterized by high concentrations of heavy metals including chromium, nickel, and cobalt, high concentrations of magnesium and iron, and low concentrations of essential plant nutrients (Safford, Viers and Harrison 2005). Combined, such characteristics contribute to poor plant productivity and high rates of plant endemism in serpentine soils (Brady, Kruckeberg and Bradshaw 2005). Most plants are unable to grow in serpentine soil, but there are a group of serpentine-indifferent plants that are able to grow in both serpentine and nonserpentine soils (Anacker 2014). By examining the microbial communities colonizing serpentine-indifferent plants on both serpentine and non-serpentine soils, we sought to determine the relative importance of plant identity and soil type on rhizoplane bacterial composition and role in growth and survival on serpentine soils.

We hypothesized that serpentine-indifferent plants host a distinct microbiome on nonserpentine soil and serpentine soils and that locally adapted serpentine microorganisms influence plant growth on serpentine soil. Additionally, we expected that soil chemistry contributes to a greater portion of variation in the root microbiome relative to plant species. We used a field survey and lathhouse experiment to answer the following questions: (1) Do serpentine-indifferent plants associate with similar microbial communities on serpentine or nonserpentine soil? (2) Is plant identity or soil chemistry the greater source of variation for soil rhizoplane bacterial communities? (3) Do serpentine microbes influence plant survival or growth on serpentine soil?

## Materials and Methods

### Study system

#### McLaughlin Natural Reserve and Hopland Research and Extension Center

Field samples were collected from serpentine and nonserpentine sites at McLaughlin Natural Reserve (McLaughlin) and Hopland Research and Extension Center (Hopland) in late March and early April of 2017. Both sites are characterized by a Mediterranean climate and hot and dry summers from April to October. Serpentine soils at McLaughlin are comprised by Henneke soil series with some Montara and Okiota series. The Henneke soils generally support chaparral, while grasslands are supported by Montara and Okiota. Hopland serpentine soil in this region is typically fine-loamy, mixed, active, mesic Typic Haploxeralfs. Representative serpentine and nonserpentine soil from McLaughlin were sent to A&L Western Laboratories, Inc (Modesto, CA) for testing using the S1B and S4 test packages. The general characteristics of the serpentine and nonserpentine soils sampled in this study are presented in Supplementary Table 1. Results are reported as mean ± standard error.

**Table 1.** – Plant characteristics of serpentine-indifferent plants

| Scientific Name | Common Name | Family | Growing Season<br>(months) | Bloom | SAS | Field | LH |
| --- | --- | --- | --- | --- | --- | --- | --- |
| *C. sparsiflora* | Spinster's blue-eyed Mary | Plantaginaceae | 3 to 9 | March through June | 1.7 | X |  |
| *G. capitata* | Blue-thimble flower | Polemoniaceae | 2 to 12 | February through April | 1.6 |  | X |
| *G. tricolor* | Bird's eye gilia | Polemoniaceae | 4 to 9 | April through August | - | X |  |
| *P. erecta* | California plantain | Plantaginaceae | 4 to 12 | March through April | 1.0 | X | X |
| *T. fucatum* | Bull clover | Fabaceae | 4 to 12 | April through June | 1.3 | X |  |
| *T. willdenovii* | Tomcat clover | Fabaceae | 3 to 12 | April through March | 1.3 |  | X |

SAS = Serpentine Affinity Score; LH = Lathhouse

#### Plant systems

Plants were chosen based on their Serpentine Affinity Mean (SAM) (Safford, Viers and Harrison 2005), where plants with high SAM are typically endemic and rarely thrive outside of serpentine soil due to competition (Anacker 2014). In this study, we focused on serpentine-indifferent (SAM=1) plants *Collinsia sparsiflora, Trifolium fucatum, Gilia tricolor, Plantago erecta, Trifolium willdenovii*, and *Gilia capitata* (Table 1; Calflora 2000). All chosen plants are annual herbs and native to California.

## Rhizoplane survey

### Plant collection

*Plantago erecta, T. fucatum, C. sparsiflora* and *G. tricolor* were collected from two geographically distinct serpentine and nonserpentine sites in Northern California in spring 2017. Full sampling details are contained in Supplementary Methods 1. Briefly, plots at each site were chosen based on the presence and co-occurrence of serpentine-indifferent plant species, with each plant species sampled from at least 2 plots (Supplementary Figure 1 and Supplementary Table 1). Plants were collected from serpentine and nonserpentine sites by excavating the whole plant, placing it into a 50-mL centrifuge tube and immediately putting the tube on ice. A garden trowel was used for sample collection and cleaned thoroughly with 70% ethanol between collections. Samples were stored at -20°C until processing and DNA extraction, outlined below.

**Figure 1.**
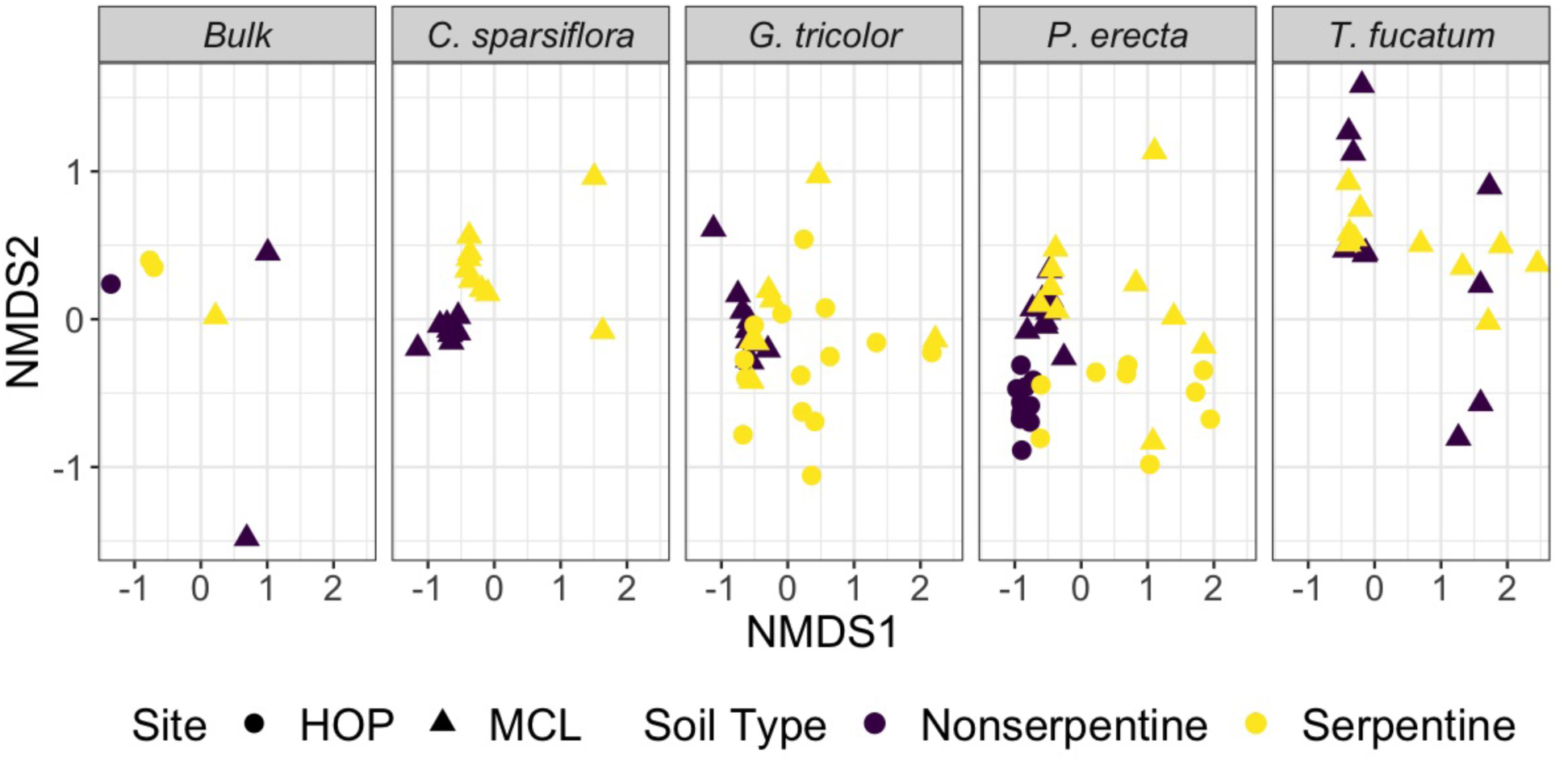
– Non-metric dimensional scaling (NMDS) plot of field-sampled bacterial rhizoplane communities associated with the plant species *Collinsia sparsiflora, Gilia tricolor, Plantago erecta*, and *Trifolium fucatum* using Bray-Curtis dissimilarity. Symbols indicate the sampling site (HOP= Hopland Reserve, MCL=McLaughlin Natural Reserve). Community composition was influenced by plant species identity (F_4,98_ = 2.97; *P*<0.001; R^2^ = 0.09), soil type (F_1,98_ = 5.33; *P*<0.001; R^2^ = 0.04) and their interaction (F_4,98_ = 1.92; *P*=0.004; R^2^=0.07).

## Lathhouse experiment

### Soil collection

Soil for the lathhouse experiment was collected from three serpentine and three nonserpentine plots at McLaughlin in late May 2016 where dense populations of *P. erecta, G. tricolor*, or *C. sparsiflora* were growing (Supplementary Figure 1). Soils within each treatment were homogenized before planting.

### Plant selection and seed sterilization

Because plant species sampled from field sites were not available for purchase and field-collected seeds germinated poorly, we used conspecifics of plants sampled in the field including *P. erecta, T. willdenovii*, and *G. capitata* grown from seeds purchased from S&S Seeds (Carpinteria, CA). Seeds were vapor sterilized for 20 hours (Clough and Bent 1998), then refrigerated until germination.

### Lathhouse soil treatments

To examine if serpentine microorganisms contribute to plant growth and survival on serpentine soils, plants were grown individually in one of four soil treatments: 1) live serpentine (serpentine), 2) autoclaved serpentine soil (autoclaved), 3) autoclaved serpentine soil amended with a serpentine microbial slurry (SS), 4) autoclaved serpentine soil amended with a nonserpentine microbial slurry (NSS). Slurries were prepared by extracting two gallons of serpentine and nonserpentine soils separately with 4L of autoclaved deionized water. Before planting, slurry solutions were added to autoclaved serpentine soil and thoroughly mixed in a large plastic tub. Tubs containing the slurry treatments were allowed to incubate at room temperature for one week. Live serpentine soil collected from the field was placed in a third tub and autoclaved serpentine soil from the field was placed in the fourth and final tub. The soil for the autoclave treatment sat overnight after initially being sterilized, then was autoclaved a second time (*Autoclave Operation and Performance Testing*; Dilly *et al*. 2004; Maignien *et al*. 2014).

### Lathhouse setup

Soil treatments were added to small D16 deepots that had been plugged with a paper towel, with each treatment replicated 24 times for a total of 288 pots (3 plant species x 4 soil treatments x 24 replicates). One seedling was placed on damp soil and covered with dry potting soil. Seedlings that did not grow were replaced after 7 days, up to six weeks into the experiment. Every week we recorded seedling survival and the number of leaves on each plant. Shoots and roots were harvested after 11 weeks of growth.

### Lathhouse shoot and root harvesting

Roots were harvested over clean parchment paper, with roots shaken to remove loosely adhering soil. The total length of the plant was collected by measuring from the root tip to the apical meristem. Roots and shoots were separated, then aboveground plant height and maximum root length measured. Shoots were weighed immediately to determine wet weight then dried at 55°C for at least 48 hours and dry weight measured. Roots were placed into a centrifuge tube and stored at -20°C until processing.

### Rhizoplane soil collection

Rhizoplane soil was collected from field-sampled and lathhouse-sampled roots. Briefly, roots were shaken in a 0.9% (w/v) NaCl and autoclaved water solution in either 15-ml or 25-mL tube depending on root size. Tubes were shaken on a lateral shaker at 10 strokes per minute (spm) for 90 minutes, then roots were removed, dried and massed. The roots were removed from the tubes after shaking and placed into a clean centrifuge tube of the same size as the original. To obtain the rhizoplane soil, 10 ml of the 0.1% (v/v) Tween80 in 0.9%NaCl solution was added to a 15-mL tube or 20 mL of the Tween80 solution was added to a 25-mL tube depending on the size of the root (Barillot *et al*. 2012). The tube and solution were shaken on a lateral shaker at 10 spm for 90 minutes. After shaking, the roots were removed from the tube and placed into a labeled coin envelope, dried at 55°C for at least 48 hours, then weighed. The tubes containing rhizoplane soil were centrifuged at 200 g for 10 minutes. The supernatant was discarded and the pellet used to extract DNA. DNA was extracted using ZR Soil Microbe DNA MicroPrep following manufacturer’s instructions (Zymo Research, Irvine, CA).

### Library prep and sequencing

From DNA extracts, the V4 region of the 16S SSU rRNA was amplified using primers 515F-806R (515F: 5’ - GTGCCAGCMGCCGCGGTAA - 3’; 806R: 5’ - GGACTACHVGGGTWTCTAAT -3’) (Caporaso *et al*. 2010). PCR was carried out in 25 µl reactions including 1 µl genomic DNA, 0.5 µl of each 10 µM primer, 12.5 µl of MyTaq Hot Start Red Mix (Bioline), and 10.5 µl of dH2O. PCR reactions were set up on ice to minimize non-specific amplification and primer dimerization. PCR conditions were: denaturation at 94°C for 2 min; 34 amplification cycles of 30 sec at 94°C, 30 sec at 51°C and 30 sec at 72°C; followed by a 10 sec final extension at 72°C. PCR products were visualized using gel electrophoresis and successful samples cleaned using Carboxyl-modified Sera-Mag Magnetic Speed-beads in a PEG/NaCl buffer (Rohland and Reich 2012).

Cleaned PCR products were quantified using the Qubit hs-DS-DNA kit (Invitrogen, Carlsbad CA), pooled in equimolar concentration and sent to the DNA Technologies and Expression Analysis Cores at the Genome Center at UC Davis for sequencing using the Illumina MiSeq platform (500 cycles v2 PE250). Raw sequences are available at https://www.ncbi.nlm.nih.gov/sra/SRP152892.

### Bioinformatics

Amplicon sequence variants (ASVs) from 16S rRNA amplicons were identified using DADA2 (v1.7.2) (Callahan *et al*. 2016a). Briefly, paired-end fastq files were processed by filtering and truncating forward and reverse reads at position 200. Sequences were dereplicated, merged and error-corrected. Chimeras were removed, and the taxonomy assigned using the SILVA database (v128) (Quast *et al*. 2013; Yilmaz *et al*. 2014; Glöckner *et al*. 2017). A phylogenetic tree based on 16S sequences was created using the ‘DECIPHER’ package (v2.8.1) in R to perform multi-step alignment and ‘phangorn’ (v2.4.0) to construct the tree (Schliep 2011; Wright 2016). The sequence table and taxonomy, phylogenetic tree and metadata, were combined into a phyloseq object and used for further analysis (phyloseq v1.22.3) (McMurdie and Holmes 2013; Callahan *et al*. 2016b). Using ‘phyloseq’, the mitochondria and chloroplast were removed from samples. Low-abundance samples (<1000 reads) were removed and the count data normalized. Alpha-diversity metrics were calculated using Shannon diversity and Simpson’s index.

### Statistical analysis

#### Field survey

To examine if bacterial communities differed in alpha diversity (Shannon or Simpson’s Index), we used ANOVA with soil type, plant species and their interaction as predictors. To visualize the similarity between groups, non-metric multidimensional scaling (NMDS) plots were created based on Bray-Curtis dissimilarity metrics (Bray and Curtis 1957; Kruskal 1964). To determine if soil or plant species differed in rhizoplane bacterial composition, we used the ‘adonis’ function from the vegan package in R. We examined if microbial ASVs differed among plant species and bulk soil, or between serpentine and nonserpentine soil using DESeq2 (v1.18.1) (Love, Huber and Anders 2014).

#### Lathhouse experiment

To determine the effects of soil treatments on seedling establishment, survival analysis was conducted on plant presence/absence data using a Cox proportional hazards regression model (‘coxph’) on a survival object (‘Surv’) in the survival package (v2.42.6). The time, in weeks, from plant absence (seeding failure to establish after weekly seedling addition) to plant presence (successful establishment) was used as the event. Visualization of the hazard ratio was achieved using ‘ggforest’ in the survminer package (v0.4.3).

To determine the effects of soil treatments on plant growth, each plant trait was analyzed using a general linearized model (‘glm’) with plant species and soil treatment as predictors and differences between group means were identified using likelihood ratio tests. Tukey HSD was used as a post-hoc test to identify differences among groups.

To examine if bacterial communities from the experiment differed in alpha diversity measured using Shannon or Simpson’s Index, we used ANOVA with soil type as predictors. To visualize the similarity between groups, non-metric multidimensional scaling (NMDS) plots were created based on Bray-Curtis dissimilarity metrics (Bray and Curtis 1957; Kruskal 1964). The ‘adonis’ function from the vegan package in R was used to determine if there were differences between the microbial communities between samples based on Bray-Curtis dissimilarity.

## Results

### Field survey

After quality filtering and removal of non-target sequences, we recovered 4,114,382 reads (average 15,825 per sample) that were grouped into 8903 amplicon sequence variants (Supplementary Data 1). Sampling curves within samples were saturating, indicating a robust sampling of the microbial diversity associated with individual plants.

Across both soil types, microbial communities of all serpentine-indifferent plants were dominated by Proteobacteria and Actinobacteria (Supplementary Figure 2). Serpentine rhizoplane communities were less diverse than rhizoplane communities formed on non-serpentine (Supplementary Figure 2, Shannon: *P*=0.007, Simpson: *P*=0.03). Plant species differed from each other in rhizoplane diversity (Supplementary Figure 2, Shannon: *P*=0.02, Simpson: *P*=0.16), although this depended on the diversity index used. Specifically, communities on *T. fucatum* were significantly less diverse than either *G. tricolor* and *P. erecta*.

**Figure 2.**
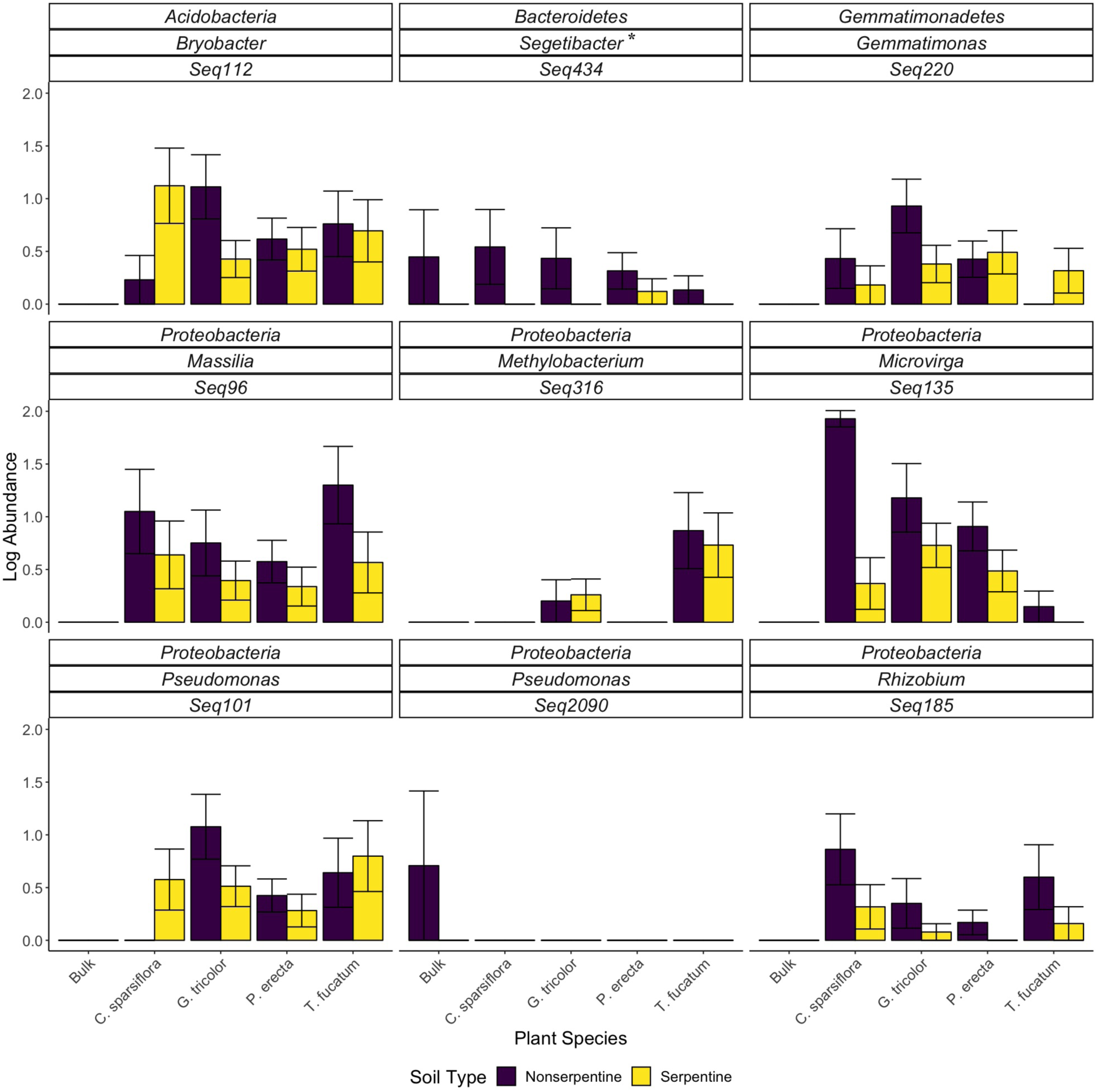
– Bacterial ASVs that were differentially abundant between plant species or between soil types from field sampling. Taxa that differed significantly between serpentine and nonserpentine soils are indicated by *, while other taxa differed between plant species at FDR <0.01 using DESeq2. Taxa shown here are representative and chosen based on their known presence on roots and in soil, role in nitrogen fixation, or presence in harsh soils. Bars indicate mean +/- SE.

Community composition was influenced by plant species identity (Figure 1; F_4,98_ = 2.97; *P*<0.001; R^2^ = 0.09), soil type (F_1,98_ = 5.33; *P*<0.001; R^2^ = 0.04) and their interaction (species x soil, F_4,98_ = 1.92; *P*=0.004; R^2^=0.07). *Trifolium fucatum* was the only plant in the study that did not associate with distinct microbial communities when grown on disparate soil types (*P*=0.13), while *C. sparsiflora* (*P*<0.001), *P. erecta* (*P*<0.001), and *G. tricolor* (*P*=0.03) all associate with distinct microbial communities when grown on serpentine or nonserpentine soil.

Differential abundance analysis showed that many of the bacteria isolated were rhizoplane-specific (Figure 2, Supplementary Data 2). Additional bacterial taxa were identified as differentially occuring among plant species. To visualize some characteristic patterns, a few ASVs were selected based on their classification as known root and soil bacteria (*Massilia, Methylobacterium, Gemmatimonas*), known role in nitrogen fixation (*Rhizobium, Microvirga*), or known presence in harsh soils (*Bryobacter*). *Pseudomonas* was chosen to demonstrate the varying association strategies of different ASVs in the same genera. *Segetibacter* was the only ASV that was differentially abundant between serpentine and nonserpentine soils (Supplementary Table 2).

### Lathhouse study

Seedling survival varied among soils with different microbial communities (*P*=0.003; Figure 3) and was typically highest in autoclaved soil and serpentine soils (live or slurry). Additionally, *P. erecta* took longer to establish than *G. capitata* while *T. willdenovii* established the more quickly than either *P. erecta* or *G. capitata*. Across all species, seedling survival was lowest in soils augmented with non-serpentine slurries.

**Figure 3.**
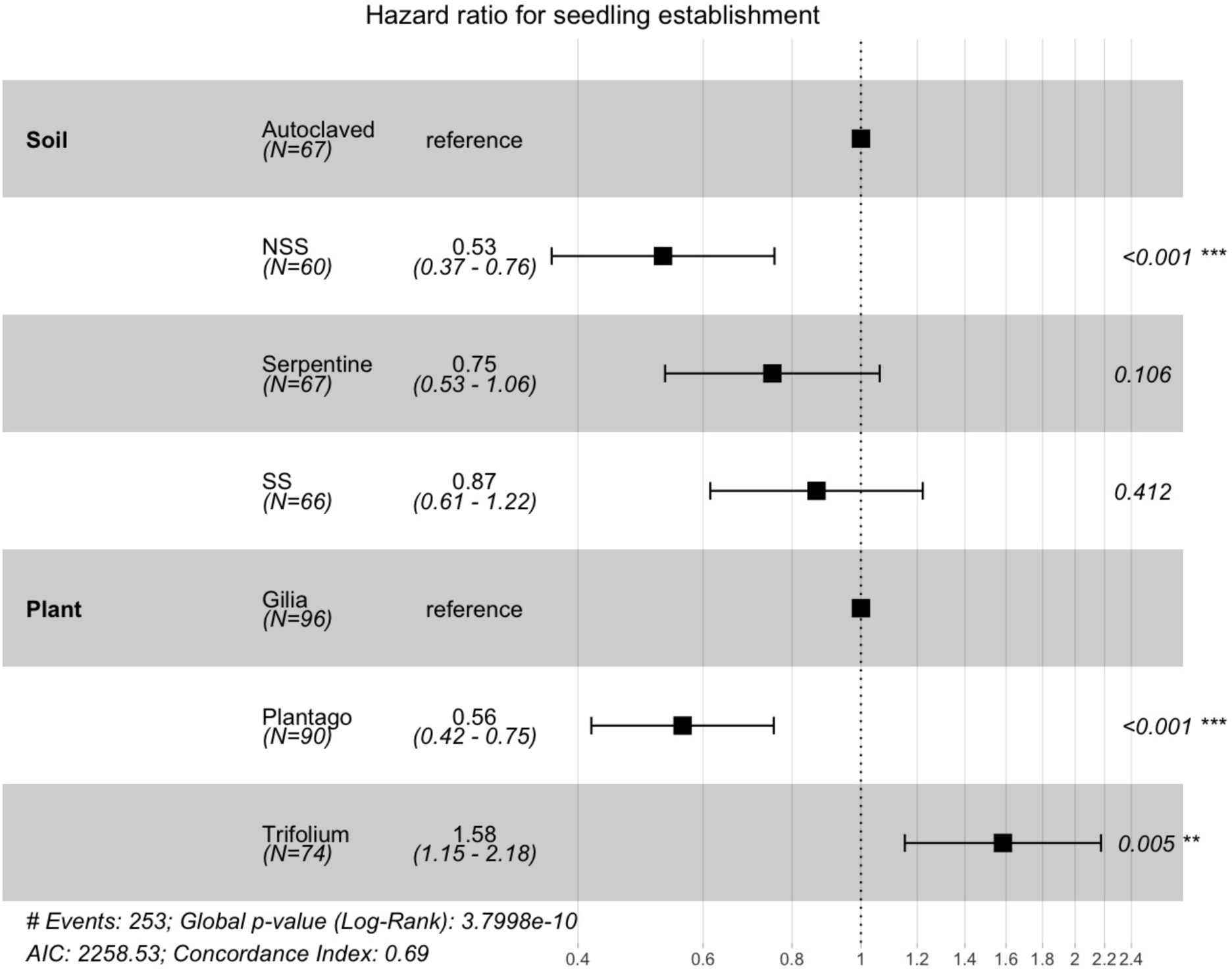
– Hazard ratio of seedling establishment in lathhouse study by soil type and plant species conducted using Cox proportional-hazards model. An HR > 1 indicates an increased likelihood of establishment while an HR < 1, on the other hand, indicates a decreased likelihood of establishment. For example, *Trifolium willdenovii* has an HR = 1.58 with a confidence interval of 1.15 – 2.18 showing that seedlings are more likely to establish relative to the reference.

Plant species varied in growth parameters and in response to soil treatments. Plant species differed in the number of leaves (Figure 4a, *X*^2^(2, N=227) = 16.56, *P*<0.001;), and although the main effect of soil treatment was not significant (*X*^2^(3, N=227) = 2.76, *P=*0.43), plant species responded differentially to soil treatments (Plant x Soil: *X*^2^(6, N=227)=12.96, *P*=0.04).

**Figure 4.**
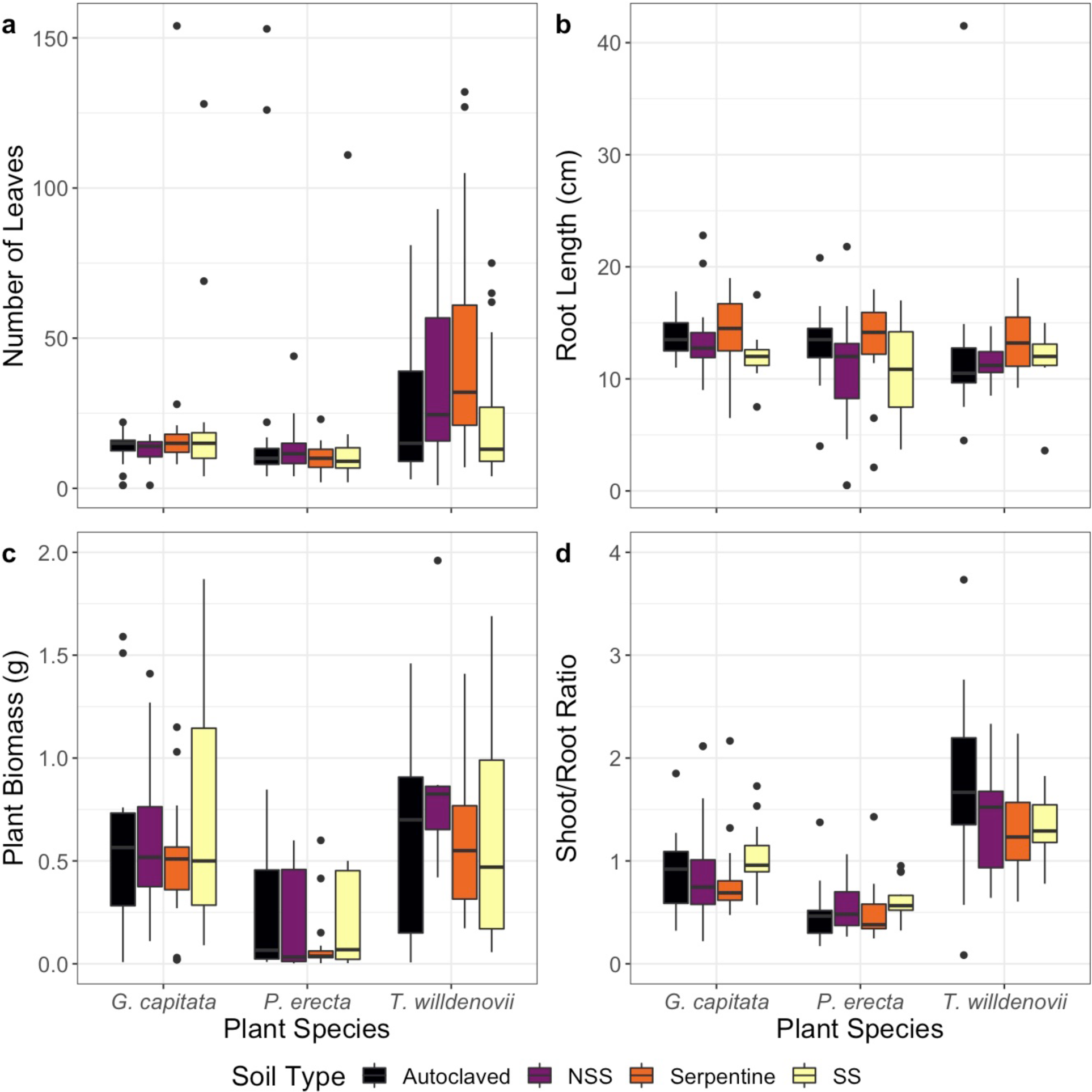
– Plant growth traits vary among plant species and soil type in lathhouse experiment. Soil type did not significantly impact (a) number of leaves (*X*^2^(2, N=227) = 16.56, *P*<0.001) (c) plant biomass (*X*^2^(2, N=193)=17.48 *P*<0.001) or (d) shoot-to-root ratio (*X*^2^(2, N=215)=9.21, *P*=0.01), but (b) root length was slightly longer in the live serpentine treatment (*X*^2^(2, N=193)=17.48 *P*<0.001).

Soil significantly influenced the root lengths of the plants (Figure 4b, *X*^2^(3, N=215)=9.95, *P=*0.02). When compared to plants grown in autoclaved soil, plant roots amended with non-serpentine slurries were, on average, 10.8% shorter, those with live serpentine soil were 2.2% longer and those amended with serpentine slurries were 15.6% shorter. Plant species varied in root length (*X*^2^(2, N=215)=6.25, *P*=0.04), but plant species did not respond differently to soils (Plant x Soil: *X*^2^(6, N=215)=1.97, *P*=0.90). Post-hoc tests did not discern differences between soil treatments.

Plant biomass varied among species (Figure 4c, *X*^2^(2, N=193)=17.48 *P<*0.001), but neither soil (*X*^2^(3, N=193)=2.85, *P=*0.42) nor species-specific responses to soils influenced plant biomass at harvest (Plant x Soil: *X*^2^(6, N=193)=6.73, *P*=0.35). Similarly, plant species differed in shoot-to-root ratio (Figure 4d, *X*^2^(2, N=215)=9.21, *P*=0.01), but neither soil (*X*^2^(3, N=215)=4.14, *P*=0.25) nor species-specific responses to soil influenced shoot height (Plant x Soil: *X*^2^(6, N=215)=9.75, *P*=0.13).

Across soil types and plant species, microbial communities in the lathhouse experiment were dominated by Proteobacteria (Supplementary Figure 3). Microbial species composition in the rhizoplane differed with soil type (Supplementary Figure 3, Shannon: *P*=0.02, Simpson: *P*=0.47) and plant species (Shannon: *P*<0.00, Simpson: *P*=0.02). In contrast to the field study, soil amended with the nonserpentine slurry were less diverse than the live serpentine soil (Shannon: *P*=0.03) and *T. willdenovii* were more diverse than *P. erecta* (Shannon: *P*<0.00, Simpson: *P*=0.01)

PERMANOVA showed that bacterial community composition differed between plant species (Supplementary Figure 4, F_3,87_ = 4.23, *P*=0.001; R^2^ = 0.09), soil treatments (F_3,87_ = 3.11, *P*=0.001; R^2^ = 0.07). There was also an interaction between soil treatment and plant species that resulted in bacterial community differences (F_9,87_ = 1.33, *P*=0.001; R^2^ = 0.09). When plant species were examined individually, both *G. capitata* (*P*<0.001) and *T. willdenovii* (*P*<0.001) associated with distinct bacterial communities depending on soil type while the community of *P. erecta* was invariant to the microbial source (*P*=0.26).

## Discussion

Taken together, our results demonstrate that both plant species identity and soil type shape bacterial species composition in the rhizoplane. Further, we show that non-serpentine microbes decrease seedling survival on serpentine soils and microbial communities differentially influence other plant phenotypic characteristics.

### Relative influence of plant identity or soil chemistry on rhizoplane bacterial communities

Similar to previous research, we found that both plant species and soil type were important in determining bacterial species composition associated with plant roots, although significant variation in species composition was explained by plot identity (13%) or remained unexplained (62%). Previous research has shown strong influence of plant species identity on root-associated bacterial community structure (Aleklett *et al*. 2015; Burns *et al*. 2015; Jorquera *et al*. 2016; Leff *et al*. 2018), however, the result documented here is particularly surprising given the large difference in physicochemical properties between serpentine and non-serpentine soils. In the field, this result was primarily driven by *T. fucatum* which associated with similar microbial communities in both serpentine and nonserpentine soils. Indeed, when *T. fucatum* is removed from the dataset, soil chemistry explains as much of the variation as plant species (soil chemistry: F_1,92_ = 6.24; *P*=0.001; R^2^ = 0.06; plant species: F_3,92_ = 2.27; *P*=0.001; R^2^ = 0.06). Nevertheless, it is notable that differentiation among plant species was still detected despite large differences in soil chemistry.

For the other serpentine-indifferent plant species including *C. sparsiflora, P. erecta*, and *G. tricolor*, bacterial species composition was strongly influenced by soil type, with communities largely distinct between serpentine and nonserpentine soil. Previous research showed *C. sparsiflora* associates with distinct fungal communities on serpentine and non-serpentine soils (Schechter and Bruns, 2008) and the current study showed that bacterial communities are distinct between soil types as well. This differentiation could be due to variation in soil properties (Supplemental Table 1) or distinct plant ecotypes found on each soil type (Wright and Stanton 2007). Ecotypic differentiation between plant populations growing on serpentine and nonserpentine soil has also been implicated in *P. erecta* (Espeland and Rice 2007). Nonserpentine and serpentine ecotypes have not been confirmed in *G. tricolor* or *T. fucatum*, but microbial community composition in *G. tricolor* may support further experimentation to determine ecotypic variation. Although ecotypic differentiation may contribute to the variation described here, the lathhouse experiment suggests that plant species identity, regardless of ecotypic variation, influences rhizoplane composition. In both the field and lathhouse studies, plant species explained more variation in rhizoplane composition than soil type, suggesting specific recruitment or growth promotion of certain bacterial taxa (e.g. Figure 2, Supplementary Data 2).

*Trifolium fucatum* did not associate with distinct communities when grown on different soil types. *Trifolium* is a genus of plants known to associate with nitrogen fixing bacteria, *Rhizobium*. Associations between *Rhizobium* and *T. fucatum* may be partially responsible for the similarity between root-association microbial communities across soil types. Still, when *Rhizobium* were removed from the data set, there was no significant difference between the microbial communities of serpentine and nonserpentine *T. fucatum* (data not shown; *P*=0.18). This may indicate the presence of other groups of microorganisms that contribute to the similarity seen across soil types, and suggests that plant species differ in the extent to which soil conditions influence their root-associated microbial communities.

Previous studies suggest that differentiation in microbial communities between serpentine and nonserpentine soils depends on the methods employed. For example, phospholipid fatty acid analysis suggested that *Avenula sulcata*, a serpentine tolerant grass does not associate with distinct bacterial or fungal communities (Fitzsimons and Miller, 2009). Similarly, 16S rRNA gene sequencing was used to show that *Acmispon wrangelianus* (Chilean bird’s foot trefoil) associated with *Mesorhizobium* when grown on serpentine or nonserpentine soil (Porter and Rice 2012). However, full genome sequencing revealed that *Mesorhizobium* isolated from serpentine harbored accessory genes, which provided a fitness advantage when grown in a high-nickel environment (Porter *et al*. 2016). Because the 16S barcoding used here may mask important functional variation in microbial communities or even populations found on each soil type, it will be important for future experiments to consider the functions or whole genomes of microorganisms in either soil environment.

### Effects of serpentine microbes on plant growth

Local adaptation of plants to soils has been studied in serpentine systems, yet the importance of locally adapted serpentine microorganisms has not received the same attention (Sambatti and Rice 2006; Wright *et al*. 2006). Here, our results suggest that microbial composition influences seedling establishment in serpentine soil, suggesting that non-serpentine microbial communities may be maladapted to enhance plant survival on serpentine. While in this experiment, measured plant traits (leaf number, plant biomass, and shoot height) were not influenced by the presence of absence of serpentine-adapted microorganisms, previous research has revealed that in some cases, microbes can influence plant growth in the presence of heavy metals. For example, *Enterobacter* sp. and *Klebsiella* sp. isolated from metal-contaminated soils produced plant-growth-promoting substances, increasing *Brassica napus* growth (Jing *et al*. 2014). Moreover, bacterial endophytes adapted to heavy metal conditions increased relative growth rates and total dry mass of *Spartina maritima* roots and leaves under these conditions (Mesa *et al*. 2015). Locally adapted ectomycorrhizal fungi improved growth of *Quercus ilex* sp. *ballota* (Holm Oak) compared to when EMF was absent in both serpentine and nonserpentine environments (Branco 2009). However, serpentine microbial communities do not always enhance the growth or survival of plant hosts. For example, serpentine arbuscular mycorrhizal (AM) and serpentine whole microbial communities decreased plant biomass relative to uninoculated plants and did not improve nickel tolerance (Doherty, Ji and Casper 2008). In our study, it is possible that different soil inoculation methods could introduce different, possibly more beneficial bacteria. In addition, it is likely that effects of microbial communities are context-dependent and may not be revealed under our experimental conditions.

Although few effects were observed on plant phenotype, soil microbial communities influenced seedling establishment, a critical barrier to survival in serpentine and nonserpentine soils. The current study did not identify specific microorganisms associated with variable establishment, but previous work has shown that pathogens can strongly mediate seeding survival (Packer and Clay 2003, Mendes *et al*. 2013). Although it is clear that the strength of plant-soil feedbacks, which can be mediated by microbial effects on seedling survival, are variable among soil types (Ehrenfeld, *et al*. 2005), it remains difficult to predict when feedbacks are important in predicting community dynamics, and if pathogens, mutualists or other microbes contribute to these effects. One promising approach may be to examine the microbial effects on seedling survivorship across soil types and mechanistic basis of local adaptation using shotgun metagenomics and whole genome sequencing of isolated bacteria in serpentine as a model system.

## Conclusions

Overall, our studies show that some serpentine-indifferent plant species associate with species-specific microorganisms on serpentine and nonserpentine soils and that those plant species showed decreased seedling establishment in the presence of nonserpentine microbes. These results suggest that both plant identity and the source microbial pool can explain variation in the structure of microbial communities, with consequences for their function. Although plant growth traits were largely indifferent to microbial community structure, the results from this experiment demonstrate that microbial communities can differ in function and suggest that serpentine microbial communities may benefit serpentine-indifferent plants on these stressful soils. Finally, these results improve our understanding of the relative influence of soil chemistry and plant identity in structuring the rhizoplane microbial community with implications for population dynamics.

## Supporting information

## Acknowledgements

We thank Cathy Koehler for assistance at McLaughlin Reserve, John Bailey at Hopland Reserve, Shenwen Gu, the Vannette Lab, and the UC Davis Data Science Initiative for their help in the data collection and analysis process as well as manuscript feedback. The UC Davis Genome center performed sequencing.

