## Supplementary material for "Plant species and soil type influence rhizosphere bacterial composition and seedling establishment on serpentine soils"

Supplementary Methods:

*Plantago erecta*, *T. fucatum*, *C. sparsiflora* and *G. tricolor* were collected from at least two serpentine and two nonserpentine sites at McLaughlin in late March, early April and early May of 2017. Five samples were collected at each site and these sites were chosen based on the co-occurrence of serpentine-indifferent plant species. *Plantago erecta* was collected from two serpentine and two nonserpentine sites at Hopland in April of 2017. Five samples were collected from each site. Fifteen samples of *Gilia tricolor* were collected from one serpentine sites at Hopland in April of 2017. Hopland sites were chosen based on soil chemistry and the presence of serpentine-indifferent plants of interest. A total of 40 plants each were collected from serpentine and nonserpentine sites at McLaughlin. At Hopland, a total of 35 plants were collected from serpentine and nonserpentine sites. Bulk soil samples were collected from at least two serpentine and two nonserpentine sites at McLaughlin and Hopland. Maps of collections sites and corresponding attribute table are available in Supplementary Figure 1 and Supplementary Table 1, respectively. Plants were collected from serpentine and nonserpentine sites by excavating the whole plant, placing it into a 50-mL centrifuge tube and immediately putting the tube on ice. A garden trowel was used for sample collection and thoroughly cleaned with 70% ethanol between collections. Samples were stored in a -20°C refrigerator until processing and DNA extraction.
