## Supplementary material for "Plant species and soil type influence rhizosphere bacterial composition and seedling establishment on serpentine soils"

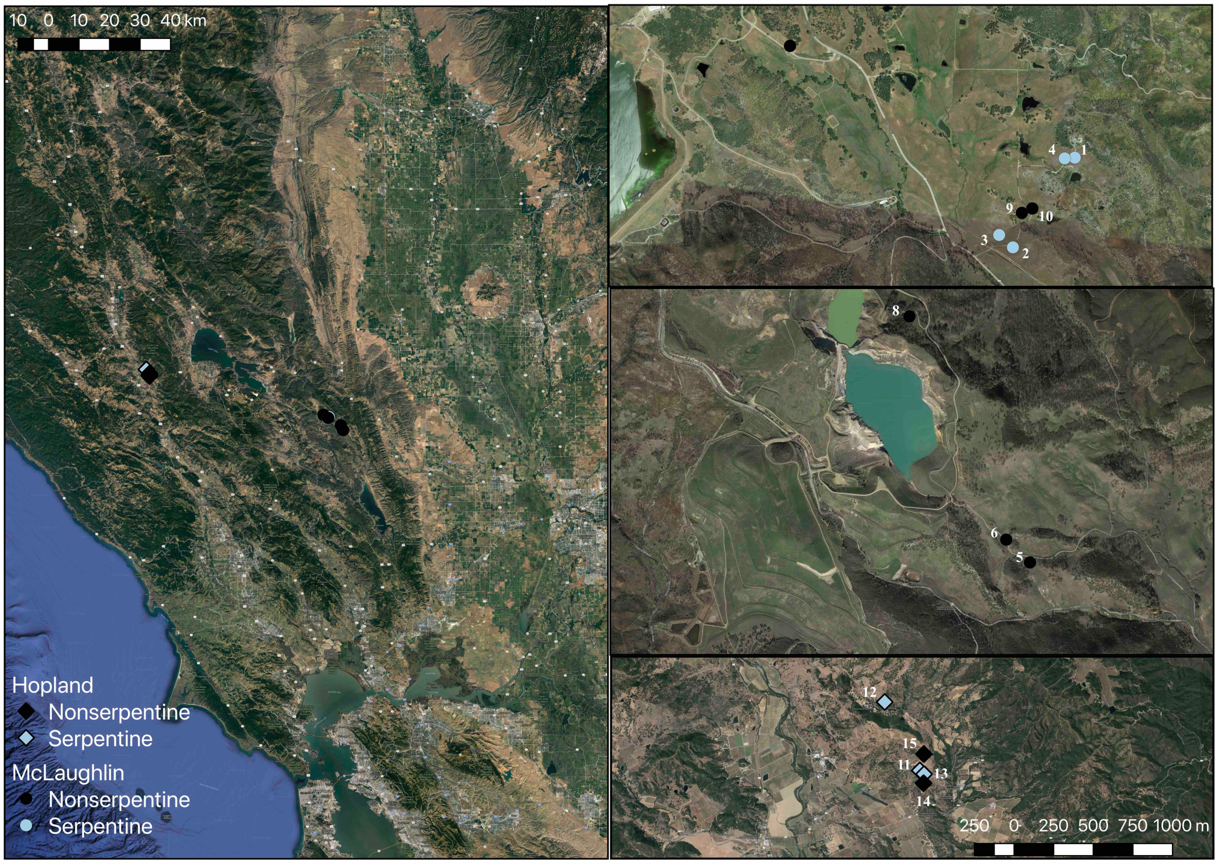


**Supplementary Figure 1** - Map of study sites at McLaughlin Natural Reserve and Hopland Research and Extension Center in CA, USA. Shapes represent sampling site (diamond = Hopland, circle = McLaughlin) and color represents soil type (blue = serpentine, black = nonserpentine).

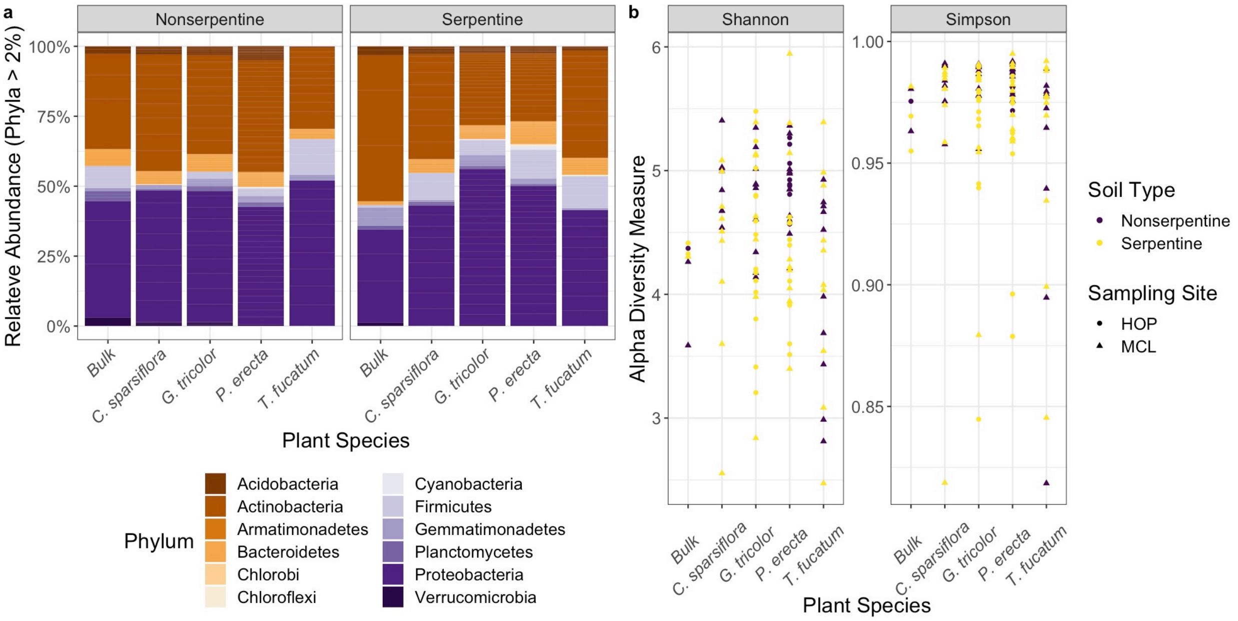


**Supplementary Figure 2** - Species composition of bacteria associated with the rhizoplane of *Collinsia sparsiflora*, *Gilia tricolor*, *Plantago erecta*, and *Trifolium fucatum*. Amplicon analysis of the 16S rRNA gene region showed that communities are predominated by (a) Proteobacteria and Actinobacteria and that (b) alpha diversity differs between plant species (Shannon: *P*=0.02, Simpson: *P*=0.16) and soil types (Shannon: *P*=0.007, Simpson: *P*=0.03).

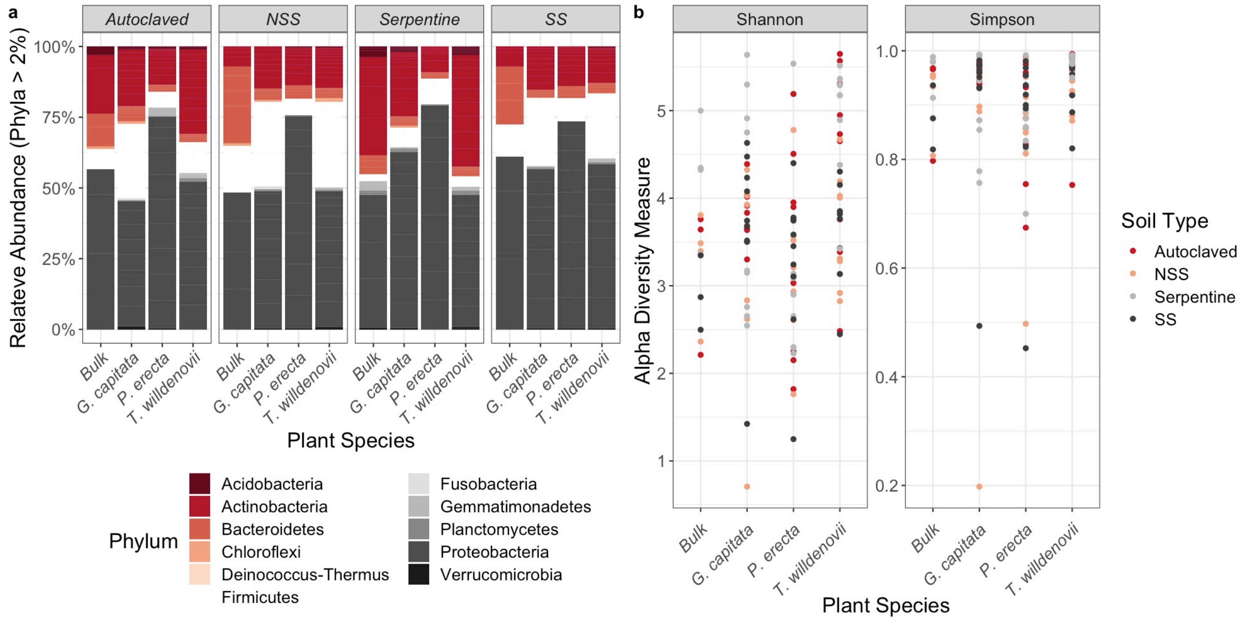


**Supplementary Figure 3** -Species composition of root-associated bacteria of *Gilia capitata*, *Plantago erecta*, and *Trifolium willdenovii*. Amplicon analysis of the 16S rRNA gene region showed that communities are predominated by (a) Proteobacteria and that (b) alpha diversity is different between soil samples (Shannon: $P$=0.02, Simpson: $P$=0.47) and plant samples (Shannon: $P$<0.00, Simpson: $P$=0.02).

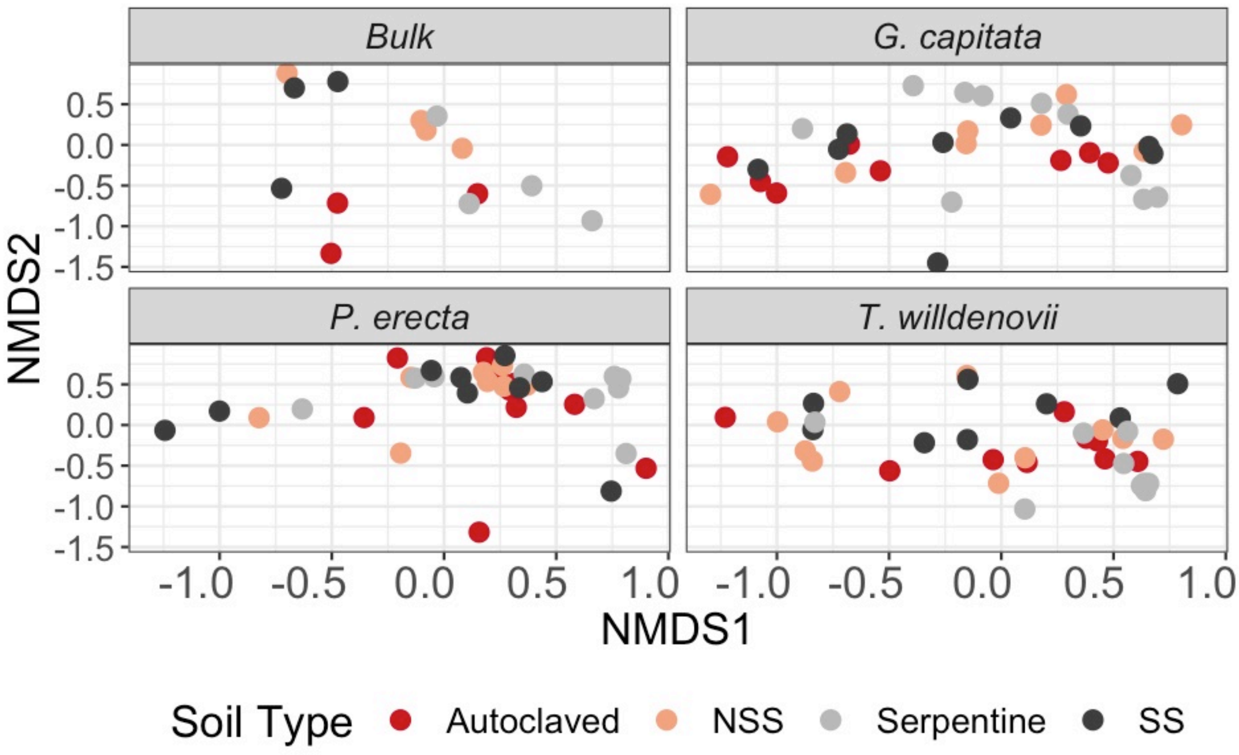


**Supplementary Figure 4** – Non-metric dimensional scaling (NMDS) plot of lathhouse-sampled bacterial rhizoplane communities associated with the plant species *Gilia capitata*, *Plantago erecta*, and *Trifolium willdenovii* using Bray-Curtis dissimilarity. Community composition was influenced by plant species ($F_{3,87}$ = 4.23, *P*=0.001; $R^2$ = 0.09), soil type ($F_{3,87}$ = 3.11, *P*=0.001; $R^2$ = 0.07), and plant-soil interactions ($F_{9,87}$ = 1.33, *P*=0.001; $R^2$ = 0.09).
